# Transcranial recording of electrophysiological neural activity in the rodent brain *in vivo* using functional photoacoustic imaging of near-infrared voltage-sensitive dye

**DOI:** 10.1101/202408

**Authors:** Jeeun Kang, Haichong K. Zhang, Shilpa D. Kadam, Julie Fedorko, Heather Valentine, Adarsha P. Malla, Ping Yan, Maged M. Harraz, Jin U. Kang, Arman Rahmim, Albert Gjedde, Leslie M. Loew, Dean F. Wong, Emad M. Boctor

## Abstract

Minimally-invasive monitoring of electrophysiological neural activities in real-time—that enables quantification of neural functions without a need for invasive craniotomy and the longer time constants of fMRI and PET—presents a very challenging yet significant task for neuroimaging. In this paper, we present *in vivo* functional PA (fPA) imaging of chemoconvulsant rat seizure model with intact scalp using a fluorescence quenching-based cyanine voltage-sensitive dye (VSD) characterized by a lipid vesicle model mimicking different levels of membrane potential variation. The framework also involves use of a near-infrared VSD delivered through the blood-brain barrier (BBB), opened by pharmacological modulation of adenosine receptor signaling. Our normalized time-frequency analysis presented *in vivo* VSD response in the seizure group significantly distinguishable from those of the control groups at sub-mm spatial resolution. Electroencephalogram (EEG) recording confirmed the changes of severity and frequency of brain activities, induced by chemoconvulsant seizures of the rat brain. The findings demonstrate that the near-infrared fPA VSD imaging is a promising tool for *in vivo* recording of brain activities through intact scalp, which would pave a way to its future translation.

## 1 Introduction

The quantification and monitoring of brain function is a major goal of neuroscience and clinical researches into the underlying mechanisms of the working brain (Friston, 2009; Raichle and Mintun, 2006). Towards this objective, several modalities have been introduced for the purpose of neuroimaging; however, existing methods have limitations. Positron emission tomography (PET) provides high molecular resolution and pharmacological specificity, but suffers from low spatial and temporal resolution (Raichle, 1998; Vanitha, 2011). Functional magnetic resonance imaging (fMRI) provides higher spatial resolution of brain activity; however, the recorded blood-oxygenation level dependent (BOLD) signal has comparatively low temporal resolution and involves uncertain interpretation (Berman et al., 2006; Logothetis, 2008). Optical imaging approaches have been used to monitor the brain function of small animals but have limited dynamic ranges and cover only superficial tissue depths because of light scattering and absorbance during penetration of biological tissue (Devor et al., 2012; Hillman, 2007). These optical approaches require invasive craniotomy for imaging of deeper brain region, with problematic long-term consequences such as dural regrowth, greater likelihood of inflammatory cascade initiation, and lack of translational practicality to non-human primate and ultimately to human studies, including those for neuropsychiatric disorders (Heo et al., 2016). Near-infrared spectroscopy (NIRS) monitors brain function non-invasively in real-time (~1ms) at several-mm depth for human brain, but suffers from poor spatial resolution (~1cm) at those depths (Strangman et al., 2013; Torricelli et al., 2014). Therefore, minimally-invasive monitoring of electrophysiological brain activities in real-time remains a task at hand in neuroimaging, with the aim to quantify brain functions in the depths of brain tissue at sub-mm spatial resolution, without need for invasive craniotomy or skull thinning techniques.

To overcome the current challenges, photoacoustic (PA) imaging has been investigated as a promising hybrid modality that provides the molecular contrast of brain function with acoustic transcranial penetration and spatial resolution (Wang and Hu, 2012; Wang et al., 2003). In PA imaging, radio-frequency (RF) acoustic pressure is generated, depending on the thermo-elastic property and light absorbance of a target illuminated by pulsed laser, and it is detected by an ultrasound transducer. Based on this mechanism, several studies have presented the capability of transcranial PA imaging (Li et al., 2018; Nie et al., 2012). Additionally, several PA approaches have been recently applied to detect electrophysiological brain activities in both tomographic and microscopic imaging configurations; Deán-Ben et al. presented *in vivo* whole brain monitoring of zebrafish using real-time PA tomography of a genetically encoded calcium indicator, GCaMP5G (Deán-Ben et al., 2016). Ruo et al. reported PA imaging *in vivo* of mouse brain responses to electrical stimulation and 4-aminopyridine-induced epileptic seizures by means of hydrophobic anions such as dipicrylamine (DPA) (Ruo et al., 2017). However, these studies used voltage sensing in the visible spectral range (488nm and 530nm for GCaMP5G; 500nm and 570nm for DPA), which may not be optimal for recording deep brain activity because of the optical attenuation. To address this, we recently presented a novel mechanism of near-infrared cyanine voltage sensitive dye (VSD) based on selective fluorescence quenching upon membrane potential variations (Zhang et al., 2017).

Here, we propose *in vivo* functional PA (fPA) imaging of chemoconvulsant rat seizure model with intact scalp using our near-infrared cyanine VSD validated by a lipid vesicle model mimicking various membrane potential levels. As a step towards minimally-invasive external neuroimaging in primates and human brains, the results demonstrate that the fPA imaging of the fluorescence quenching VSD mechanism is a promising approach to the recording brain activities of chemoconvulsant rat model at sub-mm spatial resolution, without need for any invasive craniotomy or skull thinning techniques.

## 2 Material and Methods

### 2.1 fPA VSD imaging setup

An ultrasound research system was comprised by a 128-channel ultrasound linear array transducer connected to a real-time data acquisition system (SonixDAQ and L14-5/38, Ultrasonix Corp., Canada). To induce the PA signals, pulsed laser light was generated by a second-harmonic (532nm) Nd:YAG laser pumping an optical parametric oscillator (OPO) system (Phocus Inline, Opotek Inc., USA). The tunable range of the laser system was 690-900nm and the maximum pulse repetition frequency was 20Hz. The laser pulse was fed into a fiber optic bundle delivering to bifurcated outlets, each 40mm long and 0.88mm wide (Fig. 1a). The customized, 3-D printed shell fixes the ultrasound probe between the outlets of the bifurcated fiber optic bundle outlets for evenly distributed laser energy density in lateral direction. The bifurcated output beams were overlapped at 20mm depth. The PA probe was located at around 2.2mm from bregma to obtain the cross-section of motor cortexes (Fig. 1c). The distance between fPA probe and rat skin surface was 20 mm, and the resultant energy density was at 3.5mJ/cm^2^, which is far below the maximum permissible exposure (MPE) of skin to laser radiation by the ANSI safety standards. A wavelength of 790nm was used, at which the light energy was sufficiently absorbed by the near-infrared VSD, i.e., IR780 perchlorate. Also, probing at this wavelength prevented the undesired error by time-variant oxygen change, being at the isosbestic point of Hb and HbO_2_ absorption spectra (Fig. 1b). The bias from blood context would be removed by the proposed back-end signal processing (see Section 2.9). Spatial resolution was 479.5 ± 2.7µm and 470.8 ± 24.0µm in axial and lateral directions by applying an effective signal bandwidth at 1-5MHz for envelope detection – The spatial resolution is optimized to detect the clusters of contrast chance generated by VSD redistribution in brain circuitries at sub-mm scale, rather than differentiating individual neuronal cells (~tens of µm) or micro-vasculatures (mostly ~20µm in diameter, Zhang et al., 2014a) in rat brain. See the section 2.8, criteria for selecting brain region-of-interest, and the supplementary information for detailed *in vivo* imaging performance. Fig. 1c presents a representative cross-sectional PA image of a rat brain. The dotted white outlines for the brain and motor cortex were drawn based on the rat brain atlas (Paxinos and Watson, 2014).

**Fig. 1.**
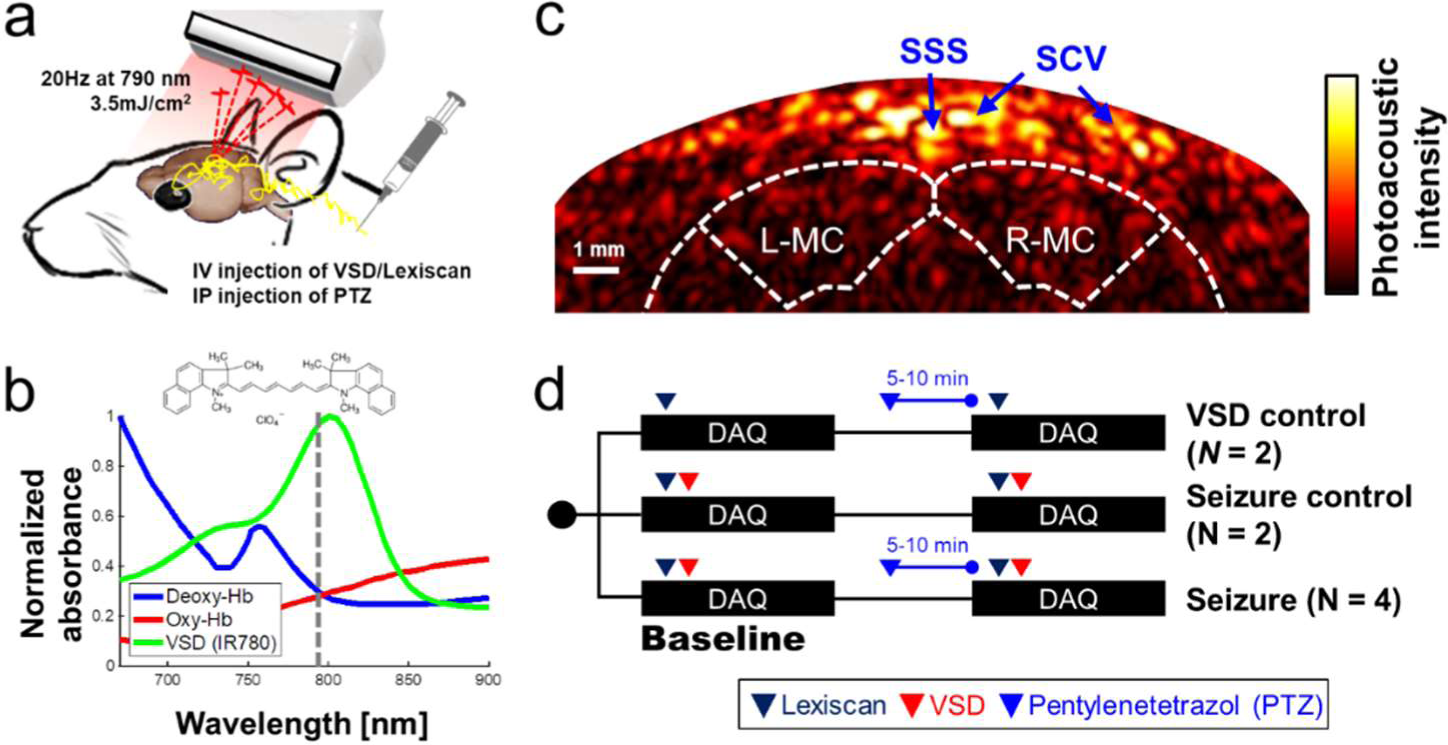
Transcranial VSD sensing setup using fPA imaging system: (a) schematic diagram of experimental setup; (b) absorbance spectra of VSD, deoxy- and oxy-hemoglobin. Dotted line indicates the wavelength used in *in vivo* experiment, i.e., 790nm; (c) cross-sectional PA image of cerebral cortex; (d) *in vivo* experimental protocol. SSS: Superior sagittal sinus; SCV: Superior cortical veins. L-MC/R-MC: left/right motor cortex. Note that the outlines for brain and motor cortex in Fig. 1c was drawn based on the rat brain atlas (Bregma 2.2mm) (Paxinos and Watson, 2014). The success of seizure induction on motor cortex was confirmed by tonic-clonic movements in the fore and hind-limbs of the anesthetized rat during the experiments (See Movie 1).

### 2.2 Fluorescence quenching-based near-infrared voltage-sensitive dye

Several cyanine VSDs have been proposed as markers for real-time electrical signal detection (Treger et al., 2014) and fluorescence tracking of electrical signal propagation on a heart (Martišienė et al., 2016). Recently we presented the mechanism of action of a cyanine VSD for fPA neuroimaging at near-infrared wavelength essential for deep transcranial neuroimaging (Zhang et al., 2017). The discussed VSD redistribution mechanism proposes a suppressive PA contrast as a product of fluorescence quenching when neuronal depolarization occurs. In the present study, we used the near-infrared cyanine VSD, IR780 perchlorate (576409, Sigma-Aldrich Co. LLC, MO, United States) with the analogous chemical structure of PAVSD800-2 in our previous study. Note that the response time of the given VSD redistribution mechanism through cell membrane should be in sub-second scale as presented in our previous study (Zhang et al., 2017).

### 2.3 Lipid vesicle phantom preparation for VSD validation

The physicochemical and biophysical studies regarding the interaction of exogenous molecules with a biological membrane have been extensively investigated using a single cell patch clamping (Jurkat-Rott and Lehmann-Hom, 2004) or using lipid vesicle models for precise control and measurement of cell membrane potential (Mazur et al., 2017; Paxton et al., 2017; Rosilio, 2018). In this study, we employ the lipid vesicle model considering our VSD redistribution mechanism in a tissue-scale between extracellular space and the cytoplasm of polarized cells as a function of membrane potential variation. The single cell patch clamping is optimized for an individual cell, rather than the cluster of cells. There is also an approach to control a cell membrane potential using valinomycin and varying the external K^+^ on a cell suspension, while being monitored in fluorescence or PA microscopy (Ruo et al., 2017). However, this treatment for membrane potential control may affect the biological state of living cells by causing them to swell and bleb in practice (Ramnath et al., 1992; Takahashi et al., 1995). On the contrary, the lipid vesicle model is free from these concerns, and more consistent measurements could be allowed between lipid vesicle model and translational *in vivo* experiments by using our cross-sectional imaging system described in section 2.1.

The lipid vesicle model was prepared using the same procedure as in Zhang et al (Zhang et al., 2017); 25-mg soybean phosphatidyl-choline (type II) suspended in 1mL of K^+^ buffer was used as the lipid vesicles. This vesicle contains 100mM K_2_SO_4_ and 20mM HEPES. The suspension was vortexed for 10 min, and followed by 60 min of sonication within bath-type sonicator to yield a translucent vesicle suspension. A Na+ buffer was also prepared, containing 100mM Na_2_SO_4_ and 20mM HEPES. Afterwards, 25:1, 50:1, and 100∶1 K^+^ gradients across vesicle membrane were established with 2.5μL, 5.0μL, and 10.0μL of lipid vesicle suspensions respectively added to 1mL of Na^+^ buffers. In the lipid vesicle model prepared, negative membrane potential (polarized state) was mimicked by adding 2.5μL of 10μM valinomycin—a K^+^ specific ionophore, thereby K^+^ ions were transported from inside to outside of vesicle membranes. On the other hand, 2.5μL of 1mM gramicidin, a nonspecific monovalent cation ionophore, enabled Na^+^ cations to move from outside to inside of vesicle membranes to short circuit the membrane potential (depolarized state). From these controls, our near-infrared VSD positively charged can move in and out through the membrane, leading to the change in fluorescence quenching depending on their aggregation status. From this lipid vesicle model, we expected the logarithmic change in membrane potential levels based on the Nernst equation (Archer, 1989): −83mV, −102mV, and −120mV. This will yield a corresponding suppression in PA intensity. The quantum yields of the VSD in depolarized states (Φ′_*F*_) were estimated based on the equations in our previous literature (Eqs. 8 and 9 in (Zhang et al., 2017)).

### 2.4 Animal preparation

For the proposed *in vivo* experiments. 8-9-week-old male Sprague Dawley rats weighing 275-390g were used (Charles Rivers Laboratory, Inc., MA, United States). The use of animals for the proposed *in vivo* protocol was approved by the Institutional Research Board Committee of Johns Hopkins Medical Institute (RA16M225). All animals were anesthetized by intraperitoneal injection with a ketamine (100mg/ml) / xylazine (20mg/ml) cocktail. (3:1 ratio based on body weight at 1ml/kg). The hair was shaved from the scalp of each rat for better optical and acoustic coupling. The head of the anesthetized rat was fixed to a stable position using a standard stereotaxic device. This fixation procedure was required to prevent any unpredictable movement during the fPA recording.

### 2.5 Chemoconvulsant seizure induction

Penetylenetetrazole (PTZ), a gamma-aminobutyric acid (GABA) A receptor antagonist was used to induce acute seizures in the animals (Löscher, 2017). PTZ suppresses the inhibitory effects of GABA, thus leading to generation of synchronized depolarizations of neurons in form of epileptiform discharges and seizures (Bradford, 1995). To induce global episodic acute seizures in rat brain, an intraperitoneal (IP) injection of PTZ (45mg/ml) was utilized based on the animal’s body weight in a volume of 1ml/kg. Subsequent doses were given if no acute motor seizure was observed in 5-10 min after the first PTZ injection. Generally, 1-2 doses were sufficient to induce the motor seizures in our experiments.

### 2.6 Pharmacological treatment for VSD delivery into blood-brain-barrier

The lumen of the brain microvasculature consists of brain endothelial cells, and the blood-brain barrier (BBB) is comprised of their tight junctions to control the chemical exchange between neural cells and cerebral nervous system (CNS). In this study, the penetration through BBB were achieved with a pharmacological method using FDA-approved regadenoson (Lexiscan, Astellas Pharma US, Inc. IL, United States). This modulates the Adenosine receptor signaling at BBB layer (Carman et al., 2011). The dosage and IV administration method indicated by the manufacturer was utilized: A volume of 150µl of the standard concentration of 0.08mg/1ml was given to each animal regardless of the weight, followed by 150µl flush of 0.9% sodium chloride for injection. The experimental protocol was designed based on the pharmacological assumption that the VSD delivery through BBB would occur during the Lexiscan’s biological half-life, i.e., 2-3 min. The efficiency of the pharmacological BBB opening was evaluated by the frozen-section histopathological analysis with near-infrared fluorescence microscopy. Three different groups were compared in this study: (1) Negative control: VSD-/Lexiscan-; (2) Control: VSD+/Lexiscan-; (3) BBB opening: VSD+/Lexiscan+.

### 2.7 *In vivo* experimental protocol

Fig. 1d shows the detailed protocol for VSD control, seizure control, and seizure groups. Note that each data acquisition was performed for 10 min to cover the biological half-life of Lexiscan for VSD delivery (2-3 min). Each dosing protocol of Lexiscan and VSD was as follows: Through the jugular vein catheter port located in the neck, 150µl of Lexiscan 0.4mg/5ml concentration was injected, and 300µl of VSD was subsequently administrated at 0.1mg/ml concentration, followed by 150µl of saline solution flush. The seizure control (*n* = 2) and seizure groups (*n* = 4) were designed to distinguish the chemoconvulsant effects on neural activity: both groups received VSD and Lexiscan, but only seizure group had IP injection of PTZ (45mg/ml/kg). The induction of seizure was confirmed by monitoring motor seizure, and another dose of PTZ was injected when no motor seizure was observed in 5-10 min. In particular, the success of the rat seizure model was determined by the behavioral observation to identify the tonic-clonic movements in whisker, fore and hind-limbs of the anesthetized rat. Once the seizure is developed, the behavioral seizure activity was maintained for entire time domain (0 – 10 min) in all the data sets presented in this paper. The VSD control group (*n* = 2) was designed to validate the inability of Lexiscan and PTZ to generate any bias on the quantification of fPA VSD responses. In this group, the baseline was obtained with the Lexiscan dosage, and subsequence data set was obtained during the chemoconvulsant seizure with secondary Lexiscan dosage without VSD.

### 2.8 Criteria for selecting region-of-interest

The coronal sections of interest were selected to include motor cortices at bregma 2.2mm, at which the synchronized depolarizations of neurons are confirmed by behavioral observation of motor seizure (See Movie 1). In the superficial depth, the signals from superior sagittal sinus (SSS) and superior cortical veins (SCV) are dominant as they contain abundant blood context. Skin surface was less obvious as the melanin contents, a major absorber in scalp, has low absorbance at near-infrared range (Jacques and Prahl, 2013). Since the intracerebral vasculatures in rat brain (mostly <20-μm in diameter, Zhang et al., 2014a) is narrower than the spatial resolution available, i.e., ~500μm, we used the relative position to SSS and brain atlas by Paxinos as a criteria to localize brain tissue region (Paxinos and Watson, 2014). In axial direction, there are four layers between SSS and brain tissue. The SSS is in the middle of dura mater (300μm) which is above the arachnoid (75μm), subarachnoid space (750μm), and pia mater (75μm) layers covering a rat brain (Nowak et al., 2011). Therefore, the expectable distance between SSS and brain atlas map should be 1,050μm, and motor cortex is extended at 3-4mm from the brain surface (bregma 2.2–3.2mm (Paxinos and Watson, 2014)). From the anatomical criteria, entire brain region was selected as the region-of-interest (ROI) to reject any subjective bias.

### 2.9 Normalized time-frequency analysis

Fig. 2 demonstrates the flow chart of our normalized time-frequency analysis method based on a short-time Fourier transform (STFT) to detect the suppressive VSD contrast in rat brain. The detailed task in each step is as following:

**Fig. 2.**
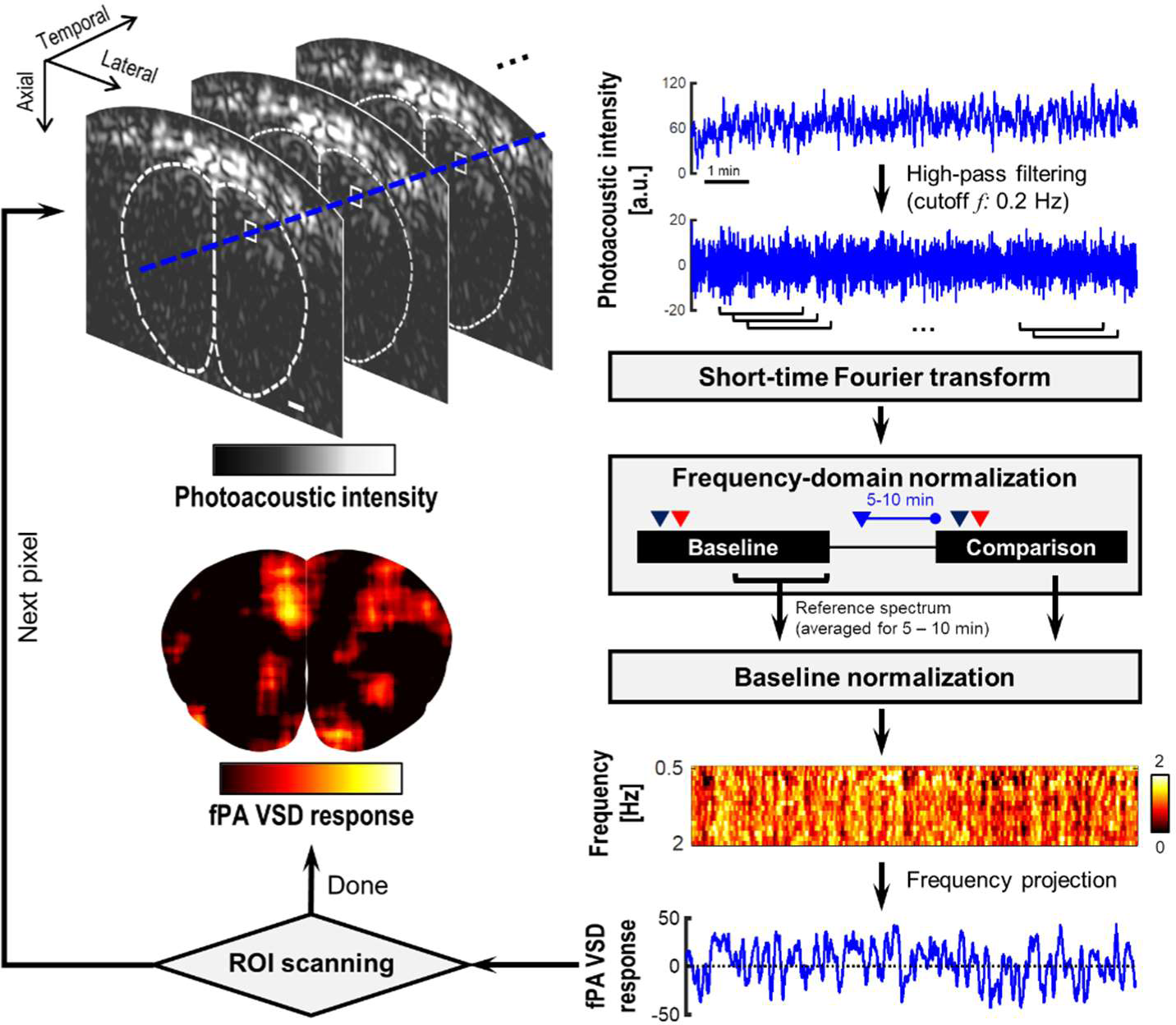
Flow chart of the short-time Fourier transform (STFT)-based normalized time-frequency analysis method. The dotted contour is boundary of brain tissue drawn based on rat brain atlas. White bar indicates 1 mm.

- **Step 1: Reconstruction of PA image sequence** using a delay-and-sum beamforming for the radio-frequency (RF) channel data averaged for 2-sec duration (40 frames) with 0.25-sec interval (5 frames interval). The signal envelope was detected in the bandwidth 1 - 5MHz. This led to 4Hz refreshing rate, and enables the frequency analysis up to 2Hz with high sensitivity. The higher imaging speed would be redundant considering slow VSD response in sub-second scale;
- **Step 2: High-pass filtering** at 0.2Hz cutoff frequency in temporal direction at each pixel of envelope-detected PA cross-section image to exclude the seizure-induced hemodynamic changes extended in few tens of seconds, which would extend up to 0.1Hz in frequency domain (Sigal et al., 2016);
- **Step 3: Short-time Fourier transform (STFT)** at each image pixel point with an analysis window across 40 temporal samples. PA(*t, f*) denotes a STFT spectrogram at a time point *t*;
- **Step 4: Frequency-domain normalization** by the averaged intensity at low-frequency band *f*_L_ (i.e., 0.3-0.5Hz) at each *t*: PA(*t, <u>f</u>*) = log10(PA(*t, f*) / *E*{PA(*t, f*)}_*f*L_), where PA(*t, f*) and PA(*t, <u>f</u>*) are the PA sequence before and after the normalization. This procedure is to fairly evaluate the amount of suppression at high-frequency range (0.5-2Hz) relative to the reference intensity at f_L_. The logarithm procedure is to present the PA intensity in negative decibel level with an emphasis on suppressive PA contrast;
- **Step 5: Baseline normalization:** PA(<u>*t, f*</u>) = PA(*t, <u>f</u>*)/PA_0_(*<u>f</u>*), in which the PA_0_(*t, <u>f</u>*) is the reference spectrum time-averaged for 5-10-min period in the baseline phase. In this step, the suppressive VSD contrast is converted into positive contrast in PA(*<u>t, f</u>*) – More suppressive contrast relative to the baseline would yield higher PA(*<u>t, f</u>*);
- **Step 6: fPA quantification of VSD response:** A fPA VSD response at each pixel is defined by a PA(<u>*t, f*</u>) projected over 0.5-2Hz range: (*E*{PA(*<u>t, f</u>*)} _*f*=0.5–2Hz_− 1) × 100. This reflects how much fractional suppression have produced compared to the reference STFT spectrum. Repeat steps 1 – 6 until all the pixels in brain cross-section are processed.

In our signal processing, high-pass filtering (Step 2) performs an important role to reject seizure-induced change in hemodynamics. All the gradual increasing bias and instantaneous changes in blood context would not be account in the outcome (Fig. 2). On the other hand, the electrophysiological seizure activity would be broadly extends from few Hz to several tens of Hz (Siemen et al., 2011), at which the suppressive VSD mechanism in fPA imaging will be presented.

### 2.10 EEG validation of neural seizure activity

To obtain the EEG records of electrical spike discharges that originated from brain tissue, sub-dermal scalp EEG recording electrodes were located at the corresponding locations on motor cortex (See the Fig. 9a), the schematic of the rat cranium (three electrodes, 1 recording and 1 reference over motor cortex, 1 ground electrode over rostrum). Silver wire sub-dermal electrodes made for use in humans (IVES EEG; Model #SWE-L25–MA, IVES EEG solutions, USA) were implanted sub-dermally, which records with a low, steady impedance, i.e., 5KΩ. Electrodes were fixed in position with cyanoacrylate adhesive (KrazyGlue, USA). The EEG signal at motor cortex was recorded with the identical preparation procedures in fPA imaging including animal preparation, administration of VSD, Lexiscan, and PTZ, time duration for recording, and interval between sequences in the protocol. Data acquisition was done using Sirenia software (Pinnacle Technologies Inc., Kansas, United States) with synchronous video capture. Briefly, the data acquisition and conditioning system had a 14-bit resolution, sampling rates of 400Hz, high pass filters of 0.5Hz and low pass filters of 60Hz. The files were stored in .EDF format and scored manually for protocol stages using real time annotations added to the recording during the experiments. EEG power for 10 sec epoch displays within the scoring software package was done using an automated module in Sirenia. Further details of our proposed EEG data acquisition and analysis used in this study are as presented in previous studies (Adler et al., 2014; Johnston et al., 2014).

## 3 Results

Fig. 3 presents the experimental results for a lipid vesicle model in various K^+^ gradient levels. The membrane potential of soybean lipid vesicle model is manipulated by the valinomycin and gramicidin, by which the quantum yield change of VSD could accordingly triggered (Fig. 3a). From the spectrophotometric and spectrofluorometric measurements (Fig. 3b), the fractional change in fluorescence emission of the depolarized state over the polarized state were 19.70%, 61.63%, and 69.69% at −83mV, −102mV, and −120mV of membrane potential levels, respectively (i.e., 25-, 50-, and 100-fold K^+^ gradient levels), while preserving a comparable level of absorbance: i.e., 0.07%, 0.70%, and 0.21% of fractional changes, respectively. The fPA intensity change presented in Fig. 3c indicates the corresponding suppressive contrast in depolarized state from a polarized state: - 4.91±4.00%, −11.49±2.00%, and −14.68±1.41% at −83mV, −102mV, and −120mV of membrane potential levels (*p* < 0.005). The expectable fPA contrast derived from the lipid vesicle experiments is −12.24±1.18% / 100 mV. The quantum yield changes according to the given K^+^ gradient levels were also estimated based on the theoretical model in our previous literature (Zhang et al., 2017). The median value in the estimated quantum yield range for each K^+^ gradient level presents a proportionally-increasing trend as depolarized (Fig. 3d). Note that the non-specific quantum yield at 25-fold K^+^ gradient is due to a limited sensitivity to differentiate the subtle membrane potential variation – The specificity of the estimation becomes proportionally improved as more K^+^ gradient is given.

**Fig. 3.**
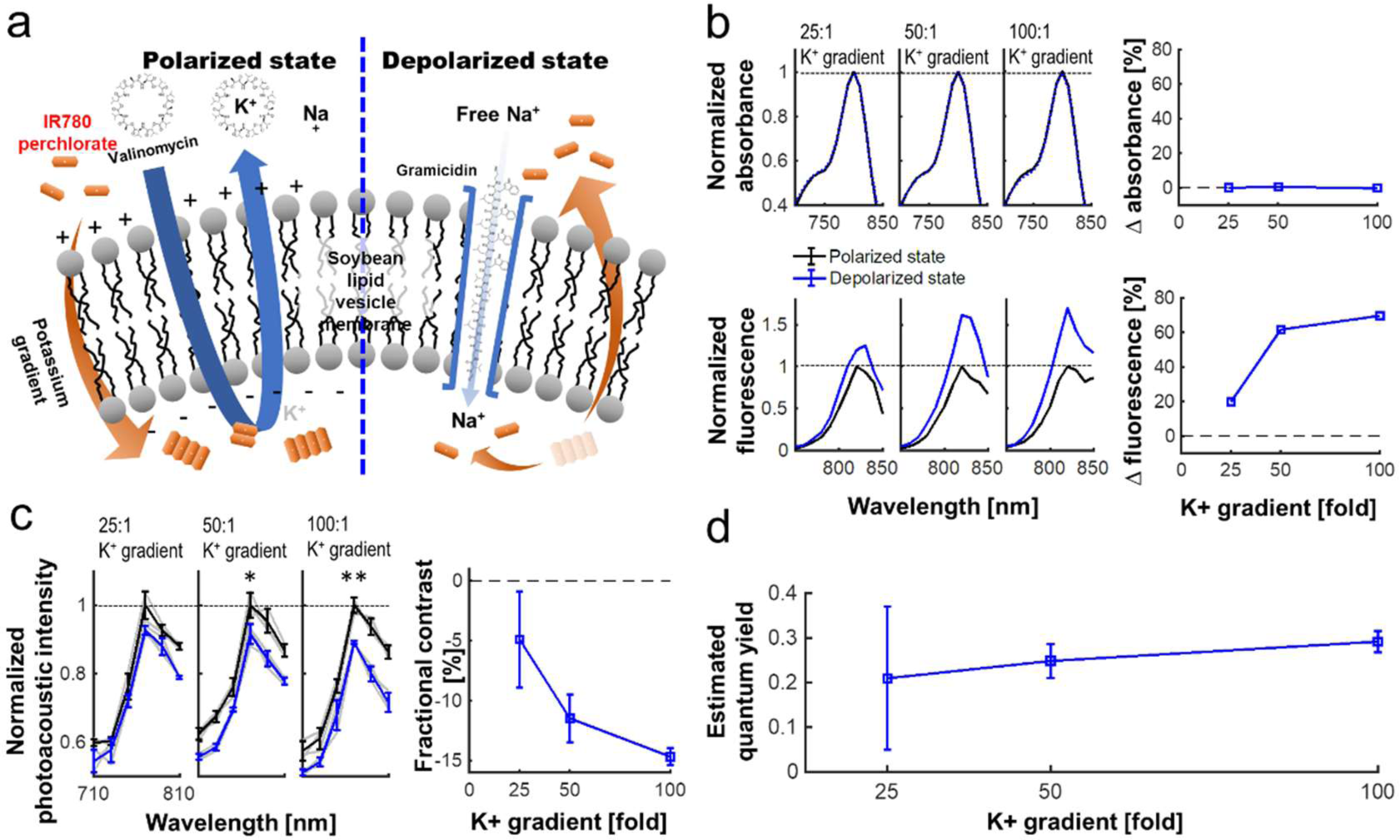
VSD characterization using a lipid vesicle model. (a) Schematic diagram of a lipid vesicle model. (b) Fractional changes of the spectrophotometric and spectrofluorometric measurements in the polarized (black) and depolarized (blue) states. (c) PA intensity spectrum at 25-, 50-, and 100-fold K^+^ gradients and fractional changes at 790 nm (*p* = 0.055, 0.010, and 0.002) between polarized and depolarized states for 25-, 50-, and 100-fold K^+^ gradients, respectively. (d) The estimated quantum yield change for each K^+^ gradient level (Zhang et al., 2017). The median values were presented in the estimated quantum yield range for each K^+^ gradient level.

With the validated VSD mechanism, we conducted the *in vivo* validation for the transcranial fPA sensing of electrophysiological neural activity in the rat brain. The fPA probe imaged the coronal cross-section at bregma 2.2mm to cover the motor cortex where the seizure was confirmed by behavioral observation. Fig. 4a shows the fPA VSD response maps projected for 10 min in each group to compare the activated brain regions among groups. All images have same range in the fPA VSD response, i.e., 0.00–3.00. In the seizure group, the chemoconvulsant seizure induced substantial VSD responses, while the control groups revealed limited activity in cortical region throughout comparison phase. Fig. 4b compares fPA VSD responses in each group projected within whole brain region. Note that they were normalized by the mean value derived from the seizure group. As a result, the seizure group scored 81.3% and 97.9% more fPA VSD response than those in VSD control and seizure control groups. The seizure group indicated significant difference in comparison to the projection of the control groups: 1.00 ± 0.31 (*n* = 4) vs. 0.53 ± 0.10 (*n* = 4); *p* < 0.05.

**Fig. 4.**
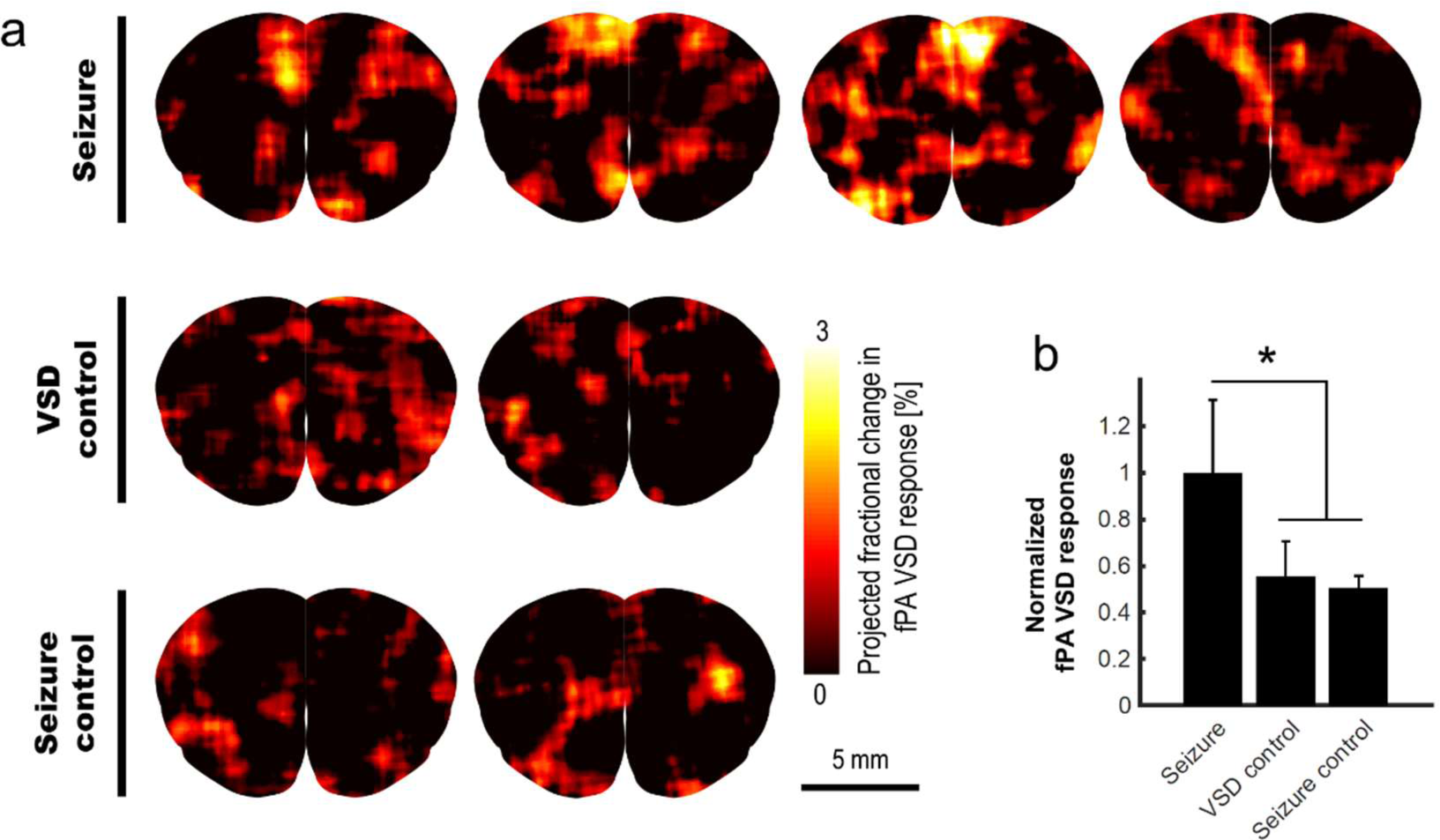
*in vivo* transcranial fPA VSD imaging for seizure, VSD control, and seizure control groups: (a) The fPA VSD response maps in each group. Note that each column indicates an individual rat included in each group; (b) The mean and standard deviation of the fPA VSD response in each group. The region-of-calculation for each rat was extended to an entire brain region. The representative examples in ROI selections and corresponding fractional PA intensity change map are presented in Fig. S2 in the supplementary information.

The appropriate VSD delivery into brain tissue was confirmed by the histopathological analysis on the harvested rat brains (Fig. 5). Three different groups were compared: (1) negative control group, VSD-/Lexiscan-; (2) control group, VSD+/Lexiscan-; and (3) BBB opening group, VSD+/Lexiscan+. From the ROIs indicated at cortical regions, the substantially-enhanced VSD uptake have been identified on the BBB opening group compared to that shown in the control group: 121.03 ± 7.14 vs. 79.19 ± 2.16; p < 0.001. The negative control group did not present any distinguishable fluorescence contrast as anticipated. The result presents the effectiveness of BBB opening based on pharmacological adenosine receptor signaling modulation by regadenoson, which is consistent with our recent fluorescence validation *in vivo*. (Pak et al., 2018)

**Fig. 5.**
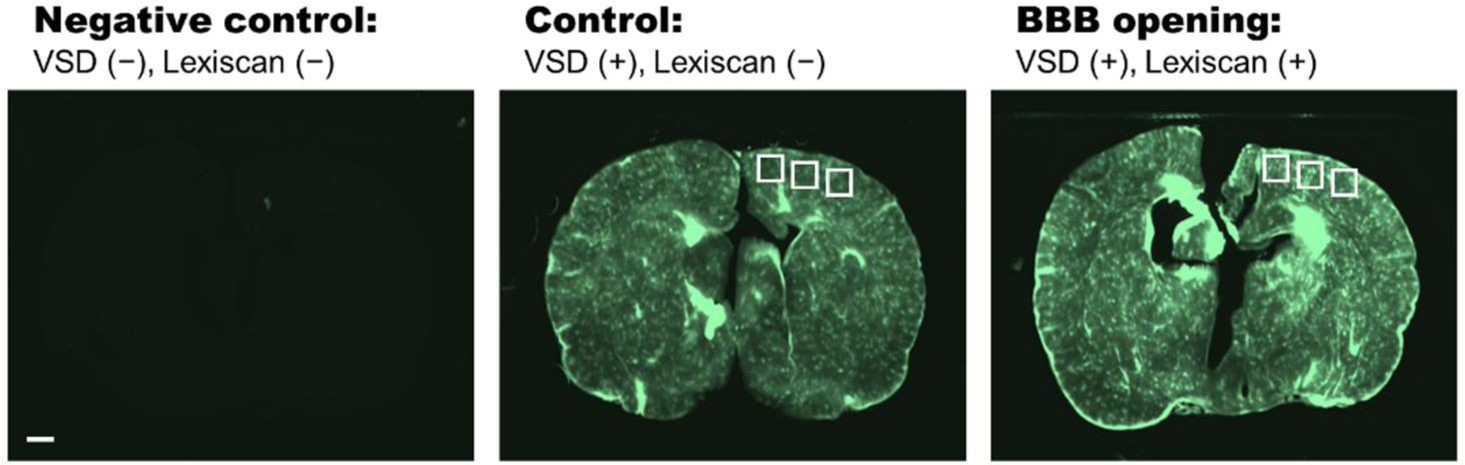
Histopathological analysis on negative control (VSD-, Lexiscan-), control (VSD+, Lexiscan-), and BBB opening (VSD+, Lexiscan+) groups. Scale bar indicate 1 mm.

We validated the chemoconvulsant-induced seizure activity in the identical *in vivo* protocol with EEG recording. Using a well-established model of chemoconvulsant-induced *status epilepticus*, we replicated the classic evolution of chemoconvulsant-induced *status epilepticus* using PTZ (Fig. 6) (Löscher, 2017). These evolutions as related to bursts of synchronized neural activity *in vivo* were assessed by EEG using the experimental protocols mirrored from that of fPA imaging experiments. We recorded vEEGs of seizure inductions using PTZ (45mg/kg IP injections) in anesthetized rats. EEG baseline recording continued until a stable seizure induction profile (i.e., continuous burst discharges indicating synchronized neuronal depolarization-related action potentials) was recorded using sub-dermal EEG scalp electrodes. The seizure activity in EEG was associated with tonic-clonic movements in the fore- and hind-limbs of the anesthetized rats, indicating motor cortex involvement (Movie 1). The PTZ evolution of status on EEG did not alter with VSD treatment.

**Fig. 6.**
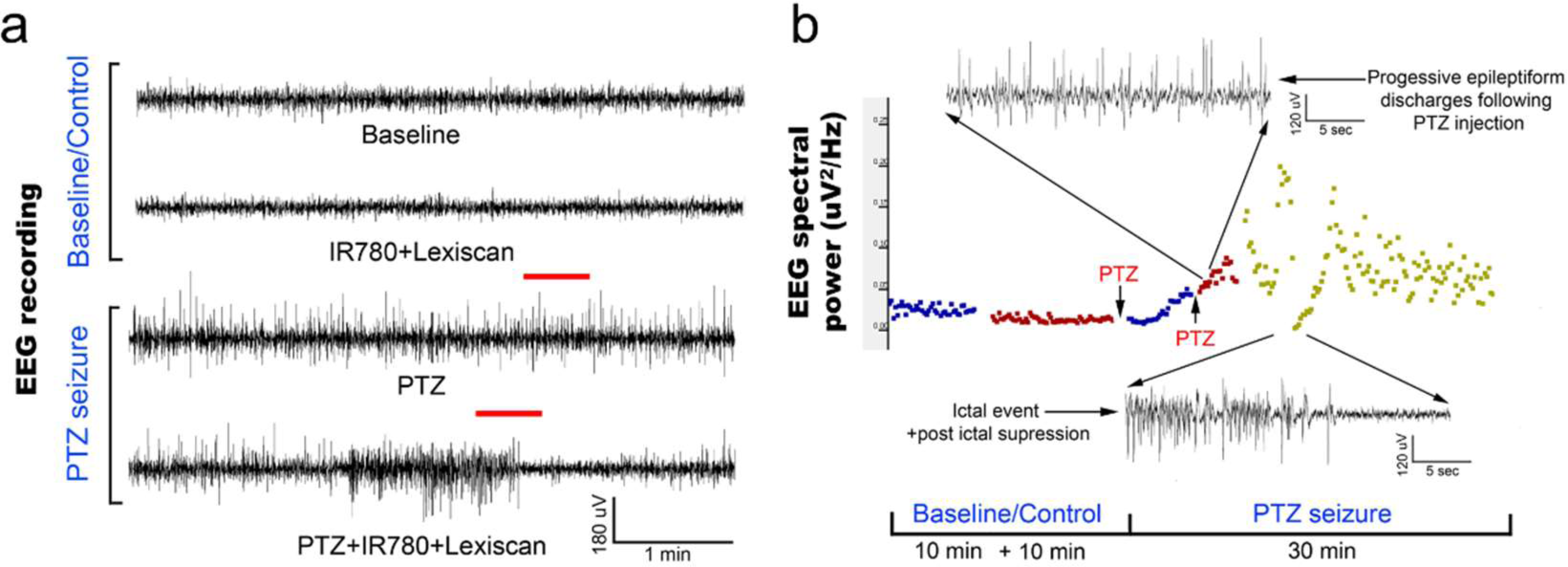
Evolution of EEG signal in the *in vivo* protocol identical to transcranial fPA imaging: (a) Representative EEG traces recorded from rat motor cortex before and during induction of status epilepticus using chemoconvulsant PTZ. The baseline and control EEG traces represent EEG activity in an anesthetized rat (see methods) with and without IR780+lexiscan given at the dosage found to not alter baseline EEG activity in the pilot study. PTS seizure induction proceeded in classical style described previously wherein episodic epileptiform burst activity evolved into status epilepticus with intermittent occurrence of seizures and stable interictal activity. (b) EEG spectral quantitation of the EEG recording done every 10 sec epoch during the EEG showed the expected progression in EEG power associated with evolution of the PTZ induced status epilepticus. Time line of PTZ injections indicated with arrows. Expanded EEG traces on top show the uniform epileptiform discharges after following second PTZ injection and below a seizure event followed of post-ictal suppression indicating the termination of that event.

## 4 Discussion

Here, we present comprehensive characterization of our near-infrared cyanine VSD mechanism using the lipid vesicle model, and a transcranial fPA VSD imaging of brain activity *in vivo* at sub-mm spatial resolution using rat seizure model with intact scalp. The near-infrared cyanine VSD, IR780 perchlorate, clearly revealed the VSD mechanism-of-action for different amount of membrane depolarization with the fractional contrast at −12.24±1.18% / 100 mV (Fig. 3). Also, the proof-of-concept *in vivo* validation study demonstrated that the non-invasive fPA VSD imaging without any invasive craniotomy or skull thinning procedures is capable of differentiating the generalized depolarization events in the seizure group from those in control groups (Fig. 4), which also well agreed with EEG validation (Fig. 6). Normalized time-frequency analysis method successfully extracted suppressive VSD contrast in coronal cross-section of rat brain over the increasing hemodynamic change with chemoconvulsant seizure using PA intensity envelope-detected in 1-5MHz bandwidth. In addition, the pharmacological enhancement of VSD delivery into rat brain by increased permeability of the BBB was confirmed by histopathological validation (Fig. 5). These results demonstrate the feasibility of transcranial fPA VSD imaging at sub-mm spatial resolution without any needs for highly-complex tomographic system and/or invasive procedures required in fPA imaging approaches at visible wavelength range.

The pixel-by-pixel correlation between fPA VSD response and the fractional PA intensity provided interesting perspectives (Fig. 7). In seizure group, the pixels presenting 2.25 to 3.00 of the fPA VSD response projected over 10 min indicated −20.94% of suppressive PA contrast, which corresponds to the proposed VSD mechanism. Interestingly, not all the suppressive changes in PA intensity was converted into high fPA VSD response, which also validate a role of the normalized time-frequency analysis method to isolate the VSD response from the hemodynamic changes. Otherwise, the control groups did not present high fPA VSD response at the cortical regions, whereas the seizure was also confirmed by behavioral observation in VSD control group. On the other hand, it would be also noteworthy that there was a case in seizure control group with unexpectedly localized yet high fPA VSD response at the primary somatosensory cortex region according to the rat brain atlas (the second rat case in Fig. 4). We hypothesize that neural activity might be real as the ketamine-xylazine does enable spontaneous and well as evoked cortical activity in anesthetized brains especially in periods after > 30 min following induction (Goss-Sampson and Kriss, 1991; Ordek et al., 2013). *In vivo* experimental protocol will be further regulated in our future investigation to reject any sensory interferences.

**Fig. 7.**
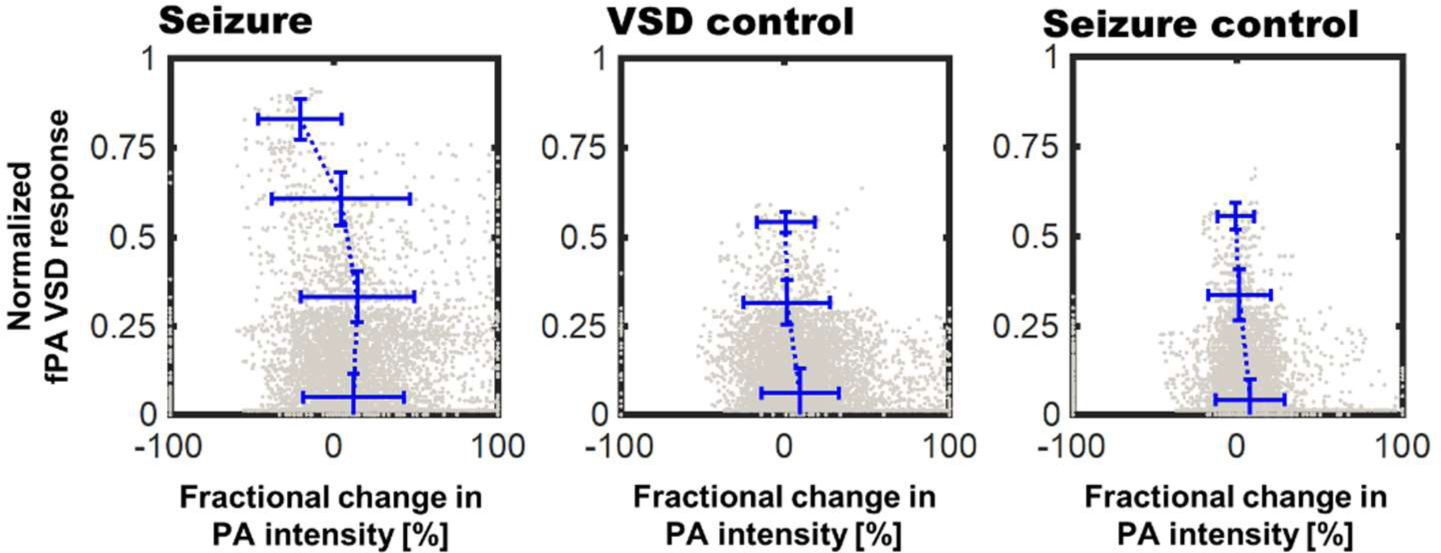
Correlation of fPA VSD response to the fractional change in PA intensity between baseline and comparison phases. Pixels were categorized to four bins of distinct fPA VSD response ranges: 0 – 0.25, 0.25 – 0.5, 0.5 – 0.75, and 0.75 – 1.

The potentially confounding factors for the *in vivo* experiments need to be carefully considered and eliminated. The change in CBV during chemoconvulsant seizure can generate proportional change of PA intensity that can be misinterpreted as the VSD response (Goldman et al., 1992; Hoshi and Tamura, 1993; Nehlig et al., 1996). Zhang et al. suggested that time frame of the CBV change induced by chemoconvulsant seizure model: The time length of gradual CBV change from PTZ injection to seizure onset was ~2 min on average (Zhang et al., 2014b). However, it was sufficiently covered by ~10 min of stabilization phase in our *in vivo* protocol (Fig. 1d). There is also an instantaneous hemodynamic change, but it extends in tens of seconds, and was rejected using high-pass filtering as described in the method section. Moreover, potential interference due to heart beating would not affect the results, as every individual fPA frame was compounded for two seconds that include 11–16 heart cycles of a rat (typically 5.5–8 beats per second).

The stability of stereotaxic fixation against the induced motor seizure was also investigated. The counter-hypothesis of this concern was an abrupt disorientation of rat brain due to motor seizure that will induce instantaneous decorrelation between adjacent PA frames. Also, based on the behavioral observation during seizure, we anticipated the decorrelation within a sub-second time scale, if it happened. For these hypotheses, we calculated the cross-correlation maps throughout PA frames obtained from 2 min to 8 min (1920 frames, 240 frames/min). Three different time intervals for decorrelation calculation were tested: 0.25 sec, 0.5 sec and 1 sec, which respectively correspond to 1, 2 and 4 frame intervals (Fig. 8). From the minimal correlation projection (MCP) map projected in entire temporal direction, motor seizure did not yield a significant decorrelation in the adjacent PA images when comparing to normal condition for the given time period. Even with 1 sec of interval, baseline and seizure phases present consistent minimal correlation value in the brain tissue region: 0.53±0.04 vs. 0.54±0.04, respectively. Therefore, the interference by motor seizure could be rejected as potential cause of artifacts in the results.

**Fig. 8.**
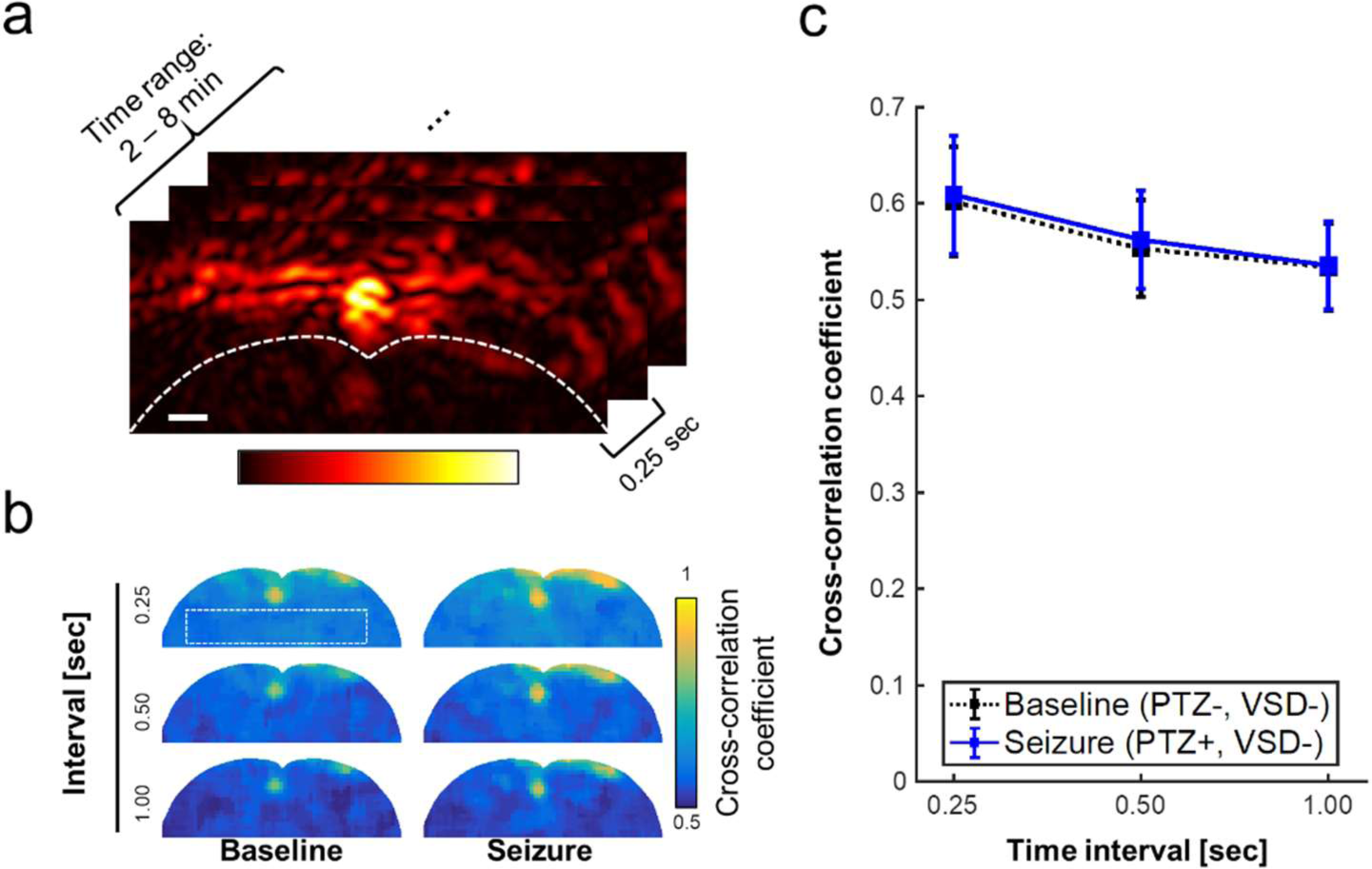
Minimal correlation projection (MCP) image using cross-correlation coefficients with varying time interval, i.e., 0.25 sec, 0.5 sec, and 1 sec, which respectively corresponds to 1, 2, 4 frame intervals with the imaging rate at 4 frames per second. (a) region of interest for the inter-frame cross-correlations, (b) MCP images of baseline (PTZ-, VSD-) and seizure groups (PTZ+, VSD-) for brain tissue region. (c) Cross-correlation coefficient for varying time intervals. Scale bar indicate 1 mm.

Toxic CNS effects of VSD is another factor that alters brain activity. We tested our protocols with varying VSD concentration in rats as a direct application to the cortex. Rats were anesthetized with IP injection to ketamine/xylazine and a cranial window was made over the right motor cortex. After recording a baseline EEG in the rat for 10-min duration with the craniotomy, the follow-on EEG recording continued to record EEG following application of increasing concentrations of vehicle alone and VSD + vehicle for the same duration of EEG recordings (i.e., 10 min) allowing comparisons of EEG responses to each increasing gradient of VSD on cortical activity as compared to baseline EEG signature in the same rat. Results for VSD with cortical application with cranial windows used in six male rats yielded reliable and reproducible EEG signatures for each concentration (Fig. 9). This protocol identified that VSD concentrations had no effect in altering the baseline EEG in the same rat, indicating no toxic effect on cortical circuit function. Direct cortical application with 100X VSD resulted in significant EEG background suppression in 4/6 rats, indicating that the certain concentrations of VSD could alter baseline circuit function in the motor cortex. This EEG suppression was recovered to baseline over the 10-min recording period, indicating that the transient effect from the time of application as the 100X VSD either diluted or cleared out of the focal application zone over the 10-min period. We reject the toxic CNS effects of VSD as we used 10X concentration based on this result.

**Fig. 9.**
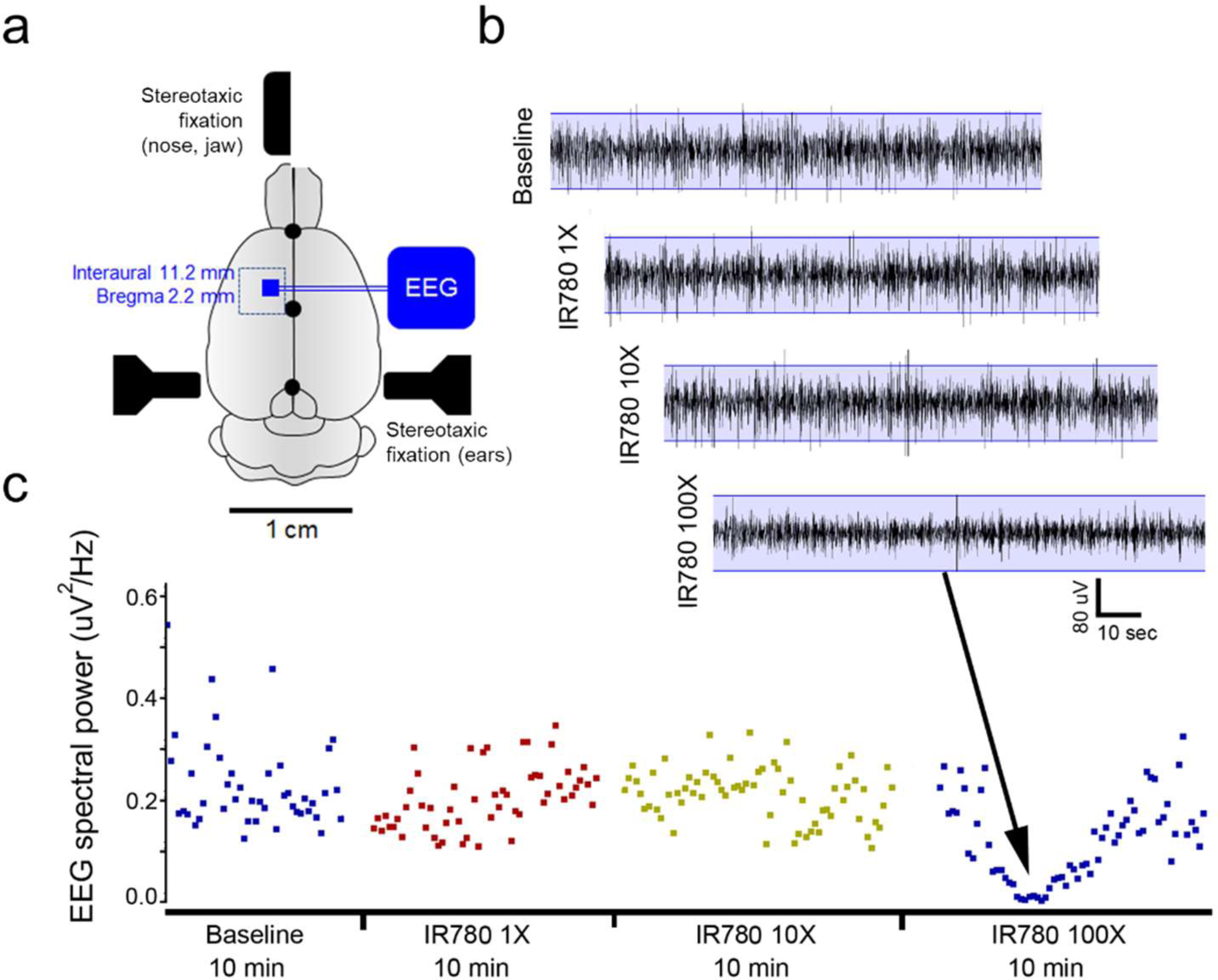
VSD toxicity study using EEG recordings during direct cortical applications using a cranial window in rats. (a) Schematic of experimental protocol. A rectangular cranial window drilled under anesthesia overlying unilateral motor cortex. Duramater was kept intact. Following craniotomy, a small window was made in duramater without traversing blood vessels. (b) EEG recording of baseline brain activity under anesthesia was followed by using a hamilton micro sssyringe to apply increasing concentrations of IR780 directly to the cortical surface via window made in duramater. Base EEG remained unaltered at lower concentrations but showed significant background suppression after applying a 100X solution. This study allowed us to determine the concentration of IR780 10X for all PA experiments. (c) EEG power spectral quantification for every 10-sec epoch of EEG over the duration of the recording confirmed EEG suppression with the 100X dose.

We plan a number of follow-up efforts to further advance the concept. We will further regulate the experimental protocol, including possible visual or audible perceptions for rats during the experiments, and also collect more *in vivo* data sets. In addition, there would be several improvements in imaging system: using 2-D PA probe would provide the most reliable setup as it enables the absolute positioning of specific brain parts. Also, the sensing speed of our current fPA imaging system would be improved. Current fPA sensing speed is limited to 4 frames per second to obtain sufficient transcranial signal sensitivity in the deep brain cortex region. This speed may limit its applicability in research, as it is well known that resting electrophysiological neural activity ranges up to several tens of Hz (e.g., delta: 1–4Hz; theta: 4–8Hz; alpha: 8–13Hz; beta: 13–30Hz; gamma: 30–80Hz). Having another dimension in spectral dimension would be beneficial to quantify the VSD response and hemodynamic changes at the same time. Successful improvements will substantially increase the capability to understand brain circuit functionality in real-time using the proposed fPA imaging technology.

To pave the way to its translation, we will further evaluate its feasibility in larger-scale brain models. Localized stimulation and detection of fPA VSD response in deeper brain regions of rodent animal have been our first step; We recently presented a success to monitor the activities in hippocampus at ~5mm depth through intact scalp in rat animal (Kang et al., 2018b). In this study, a dentate gyrus (DG) gatekeeping function was selectively stimulated by focal N-methyl-D-aspartate (NMDA) infusion using a reversed microdialysis, while collecting dialysate samples by a forward microdialysis. On the other hand, the fPA VSD neuroimaging in sagittal direction was concurrently performed at the contralateral side of the microdialysis. The configuration enabled the quantification of an extracellular glutamate concentration as a marker of excitatory neurotransmittance focally manipulated at the DG and its correlation to fPA VSD response. As a result, we presented the positive correlation of fPA VSD response to the dose-dependent changes of extracellular glutamate concentration at the hippocampal circuitry. We will also step forward to use the larger brain models of porcine and non-human primate animals, which will provide practical size and anatomy of brain as well as thicker skull and scalp when compared to humans. We also achieved an encouraging progress to obtain the sufficient sensitivity on the physiological hemodynamic changes through thick scalp and skull layers intact with 5mJ/cm^2^ energy density (Kang et al., 2018a).

Having near-infrared VSD with a faster time response is definitely in interest for our further studies, and we are working on to secure better kinetics and absorbance. It would be largely beneficial for more profound level of neuroscientific researches. For example, evaluating instantaneous responses to various controlled stimulations at a specific neural compartment would also requires the faster VSD. However, use of such faster VSD would necessitate much higher standard in fPA imaging sensitivity to overcome background noise caused by various factors such as laser energy fluctuation, heart beating, and other biological variations, etc. Therefore, faster laser system with sufficient energy density will be needed to secure a sufficient transcranial imaging sensitivity. In addition, employing advanced image quality enhancing algorithms such as deep learning-based filtering or adaptive beamforming would be also investigated.

Even though we succeed to detect brain activities at the VSD concentration below the threshold interfering brain activity (Fig 9), there have been no long-term and comprehensive toxicity study. The toxicity and biodegradability of our VSD is an important issue that deserves further evaluation. However, we are optimistic about this issue as the metabolic products of IR780 perchlorate should be very similar to ICG, FDA-approved near-infrared cyanine dye, because they are basically comprised by same chromophore. This strongly suggests its biocompatibility of our cyanine VSD. We will further prove our hypothesis in our future works.

Furthermore, the integration of localized neural stimulation methods to our fPA imaging will allow us to substantially elevate our understanding on how brain respond to a controlled stimuli (Lewis et al., 2016). The integration of the proposed fPA VSD imaging with a transcranial neuromodulation method may have a huge impact on the neuroscientific and clinical efforts by enabling the breakthrough beyond the passive brain investigation. In addition, there could be additional benefits on non-pharmacological BBB opening with a specific modality such as focused ultrasound (Chu et al., 2015; Tufail et al., 2011).

## 6 Author Contributions

J. K. planned and conducted lipid vesicle and *in vivo* experiments, analyzed the data, and wrote the first draft; H. K. Z. conducted lipid vesicle experiments and analysis; S. D. K. conducted *in vivo* EEG measurements and analysis; J. F. and H. V. prepared animal model and conducted *in vivo* experiments; P. Y. and L. M. L. provided VSD compound with the guidance for its *in vivo* use. A. P. M. and M. M. H. conducted histopathological validation of VSD delivery to brain; J. U. K. funded and participated in the development of the current fPA imaging system; D. F. W., A. R. and A. G. provided the original funding for the *in vivo* experiments; E. M. B led the development, system specification, and funding of the current fPA VSD imaging system. He also confirmed the final manuscript. All authors edited the manuscript.

## 8 Acknowledgments

Authors thank Dr. Abhinav K. Jha for proof reading the manuscript and providing helpful comments. Also, authors thank Drs. Diane S. Abou and Daniel L. J. Thorek for providing equipment and facilities for lipid vesicle experiments with helpful comments.

