## Supplementary Information for "Transcranial recording of electrophysiological neural activity in the rodent brain *in vivo* using functional photoacoustic imaging of near-infrared voltage-sensitive dye"

### 1 Characterization of fPA imaging performance.

The performance of the PA imaging system was evaluated. The wire target with 150-μm diameter was placed in the water tank, and the PA data were obtained at 20-, 25-, and 30-mm distance which are in the depth-of-interest for our lipid vesicle and *in vivo* experiments. The PA image sequence was reconstructed by using a delay-and-sum beamforming, and envelope was detected for the bandwidth 1 - 5MHz to obtain both deep imaging depth and sub-mm spatial resolution. The center frequency and fractional bandwidth of the ultrasound linear array transducer are 6.88 MHz and 78.25% at -6dB level, covering the imaging bandwidth used in this study. The data recording was also extended for 10 min to validate the expectable range of pulse-to-pulse laser energy fluctuation.

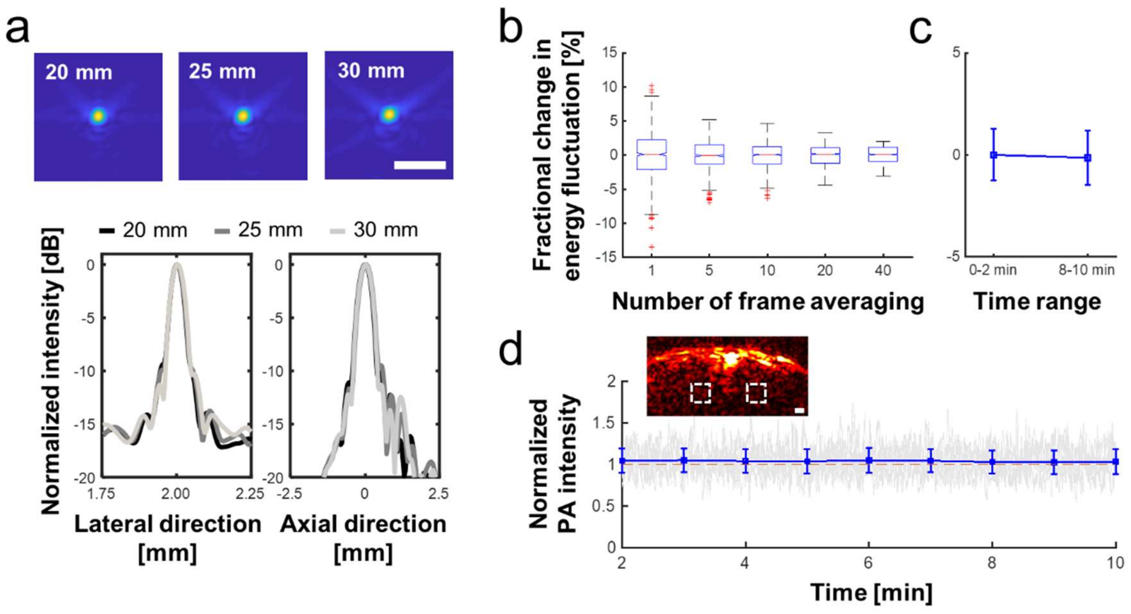


Fig. S1. PA imaging characterization: (a) spatial full-width-half-maximum (FWHM) beam profile in axial and lateral directions at various depth-of-interest (i.e., 20, 25, and 30 mm). (b) Fractional change in energy fluctuation with various number of frame averaging (i.e., 1, 5, 10, 20, 40 frames). (c) Long-term comparison in laser energy fluctuation between 0-2 min to 8-10 min time ranges. (d) Normalized PA intensity for 8 min. PA intensity averaged for 1-2 min was used as a reference intensity. White dotted rectangular in the PA image of seizure control rat indicates the photo-bleaching ROIs.

Fig. S1a presents the full-width-half-maximum (FWHM) measurements were 473.0 µm, 477.8 µm, and 482.6 µm (479.5 ± 2.7 µm in average) in axial direction, and 448.5 µm, 467.6 µm, and 496.2 µm (470.8 ± 24.0 µm in average) in lateral direction at 20 mm, 25 mm, and 30 mm depths, respectively. We also evaluated the short- and long-term laser energy variation; Fig. S1b shows the energy fluctuation when various number of frame averaging is applied: 1, 5, 10, 20, 40 frames. The result proves that the median values were well converged to baseline: 0.10%, -0.07%, 0.06%, 0.14%, and 0.10% of fractional change from the global mean value, respectively. Also, the lower and upper adjacent values were limited well with more frame averaging: -8.69 – 8.69, -5.17 – 5.13, -4.52 – 4.86, -3.30 – 4.34, and -2.01 – 3.02 for 1-, 5-, 10-, 20-, 40-frame averaging, respectively. Note that our image reconstruction was with 40-frame averaging. *In vivo* signal fluctuation was much greater than error given by laser fluctuation (raw PA intensity sequence in Fig. 2). In the investigation on long-term laser energy variation, the fractional PA intensity change was only 0.15% between the first and last two minutes: 0-2 min: 0.00±1.27% vs. 8-10 min: 0.15±1.33 % (Fig. S1c). In addition, there was no significant photo-bleaching detected with given energy density, 3.5mJ/cm$^2$ (Fig. S1d).

### 2 Selection of region-of-interest (ROI) for the quantification of fPA VSD response.

Fig. S2 presents the representative examples in ROI selection for fPA VSD response quantification based on the anatomic relationship between SSS and brain tissue. Whole brain tissue region was selected using a rat brain atlas at bregma 2.2mm as used in the *in vivo* experimental setup. The fractional change of PA intensity at 790 nm between comparison and baseline phases were also presented (second row), which demonstrate the seizure-induced CBV change. Note that the PA image was projected in last 5 min for both comparison and baseline phases.

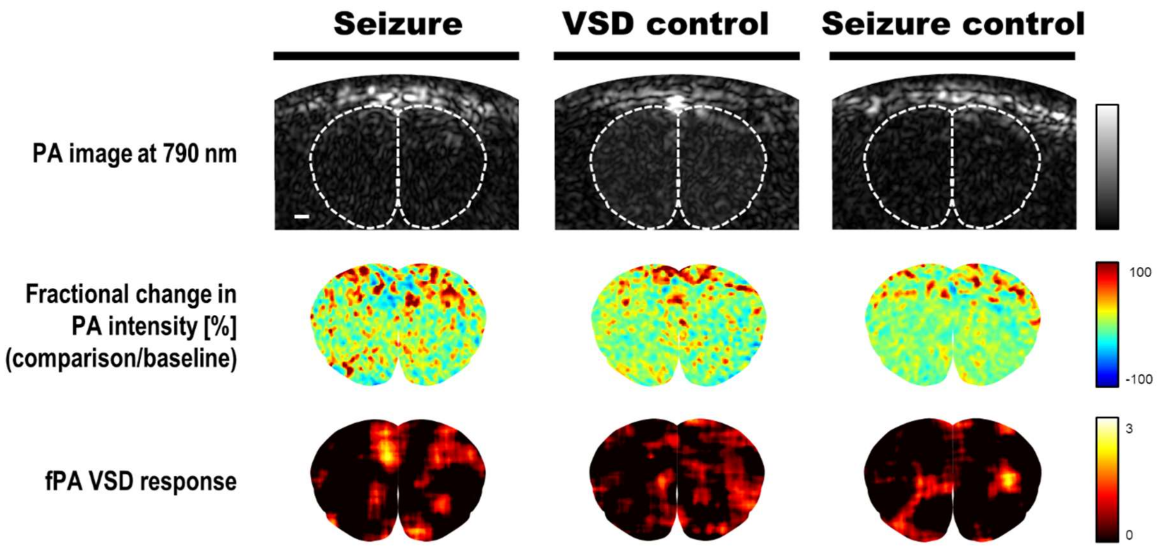


Fig. S2. Representative examples in ROI selection for fPA VSD response reconstruction and its correlation to the corresponding fractional change in PA intensity from baseline to comparison phases.
